# Angiocrine factors from HUVECs amplify erythroid cells

**DOI:** 10.1101/837823

**Authors:** Ryohichi Sugimura, Ryo Ohta, Chihiro Mori, Emi Sano, Tatsuki Sugiyama, Takashi Nagasawa, Akira Niwa, Yu-suke Torisawa, Megumu K. Saito

## Abstract

Erythropoiesis is regulated by microenvironmental factors from the vasculature. Enhanced erythropoiesis, which occurs under stress or during development, amplifies erythroid cells to meet the demand of red blood cells. This process uncouples cell division and differentiation, thus the accumulated erythroid cells remain undifferentiated in the vasculature. However, little is known about how vascular endothelial cells (ECs) regulate erythropoiesis. Here we identified that human umbilical vein endothelial cells (HUVECs) keep erythroid cells undifferentiated and amplify their number. We determined that HUVECs amplify erythroid cells via secreted angiocrine factors. The expression profile of these factors suggested that they resemble macrophage-crines for enhanced erythropoiesis. Molecularly, HUVECs mediate the activation of ERK signaling. These data indicate that angiocrine factors from HUVECs enhance erythropoiesis via the amplification of undifferentiated erythroid cells. Our study contributes to the ultimate goal of harnessing erythropoiesis to replace blood transfusions.

## Introduction

Every year, approximately 36,000 units of red blood cell transfusions are used for patients suffering from diseases, undergoing surgical operations, or some other medical need every day in the US (https://www.redcrossblood.org/donate-blood/how-to-donate/how-blood-donations-help/blood-needs-blood-supply.html). The source of these transfusions has depended on blood donors, but more stable sources are required for the anticipated growth in demand (Batta, 2016). One alternative is stem cells, which can be proliferated to high numbers that pro duce the needed volume (Chung, 2017; Doulatov, 2013; Kinney, 2019; Orkin, 2008). However, limited understanding of erythropoiesis has prevented the production of clinically relevant quantities of erythrocytes (Fang, 2016; Wei, 2019).

Erythropoiesis is the process through which red blood cells are produced (An, 2015). Erythropoiesis begins with the commitment of hematopoietic progenitor cells to erythroid cells that takes places in both embryos and adults (Nandakumar, 2016; Pimkin, 2014). Under stress erythropoiesis, glucocorticoids uncouple cell division and differentiation, thus maintaining and amplifying undifferentiated erythroid cells (Li, 2019).

The amplification of erythroid cells takes place in the vascularized regions of tissues, such as yolk sac, fetal liver, placenta, and adult bone marrow (BM) (Baron, 2012; Van Handel, 2010). The role of vascular endothelial cells (ECs) has been documented in the maintenance of hematopoietic stem cells, the differentiation to both myeloid and lymphoid lineage types, and the production of platelets (Morrison, 2014; Pinho, 2019). However, the role of vascular ECs in erythropoiesis is unclear. Elucidating the contribution of vascular ECs and their effector molecules is expected to achieve the clinically relevant number of red blood cells from stem cells (Butler, 2010; Ziyad, 2018).

Herein we demonstrate the role of human umbilical vein endothelial cells (HUVECs) in enhanced erythropoiesis. We show that angiocrinie factors secreted from HUVECs maintain and amplify undifferentiated erythroid cells. We profiled these angiocrine factors and found that they shared features with macrophage-crines known to enhance erythropoiesis (Lopez-Yrigoyen, 2019). The prospective downstream target of the angiocrine factors is ERK signaling (Eblen, 2018; Grasman, 2017; Rezaei, 2019; Smalley, 2018), whose suppression terminated the HUVEC-mediated amplification of erythroid cells.

## Results

### HUVECs amplify erythroid cells

To address whether embryonic ECs have a role in erythropoiesis, we co-cultured human BM-CD34+ hematopoietic progenitor cells with either HUVECs or human pluripotent stem cell (hPSC)-derived ECs seeded on fibronectin. Co-culture with HUVECs resulted in the amplification of Pro-erythroblasts (EB)s (CD71+GLY-A−) and EBs (CD71+GLY-A+), so that overall total of erythroid cells increased (fig. 1A-B). HUVECs on fibronectin stalled the waterfall pattern of erythroid differentiation, particularly before the entry to Late-EB (CD71−GLY-A+) stage (fig. 1A). In contrast to HUVECs, co-culture with human PSC-derived ECs fully differentiated to Late-EBs (fig. 1A). We measured the differentiation potential of the amplified erythroid cells with HUVEC co-culture. We isolated and cytospun the EB population to identify basophilic erythroids (fig. 2A). To examine if the amplified erythroid cells could undergo further differentiation, we cultured them for an additional week. We found that 32% of them exited from EB to become Late-EB and differentiated to orthochromatic erythroid (fig. 2B), indicating that differentiation capacity could be restored. These data demonstrate that HUVECs amplify erythroid cells by keeping them undifferentiated.

**Fig. 1.**
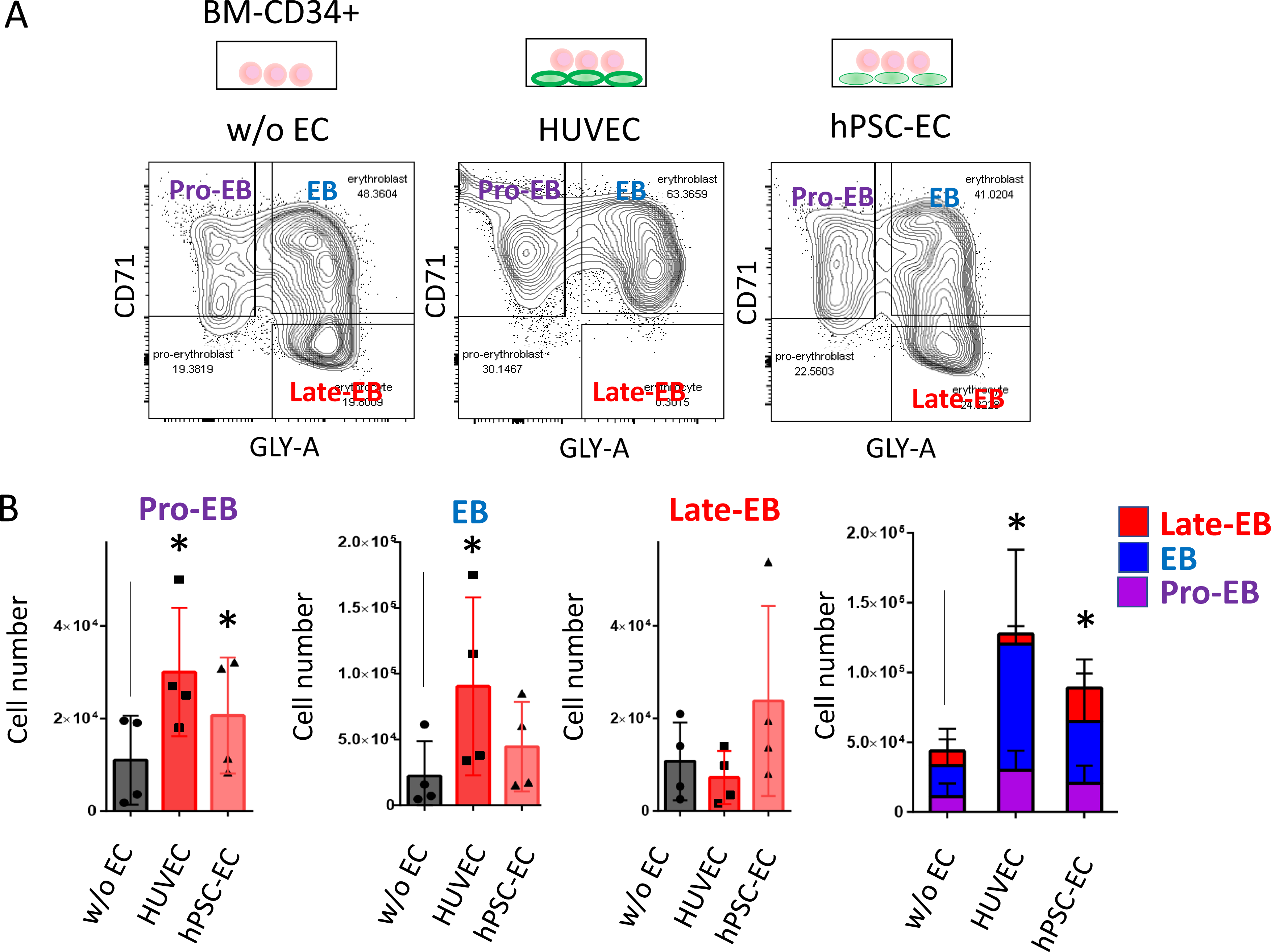
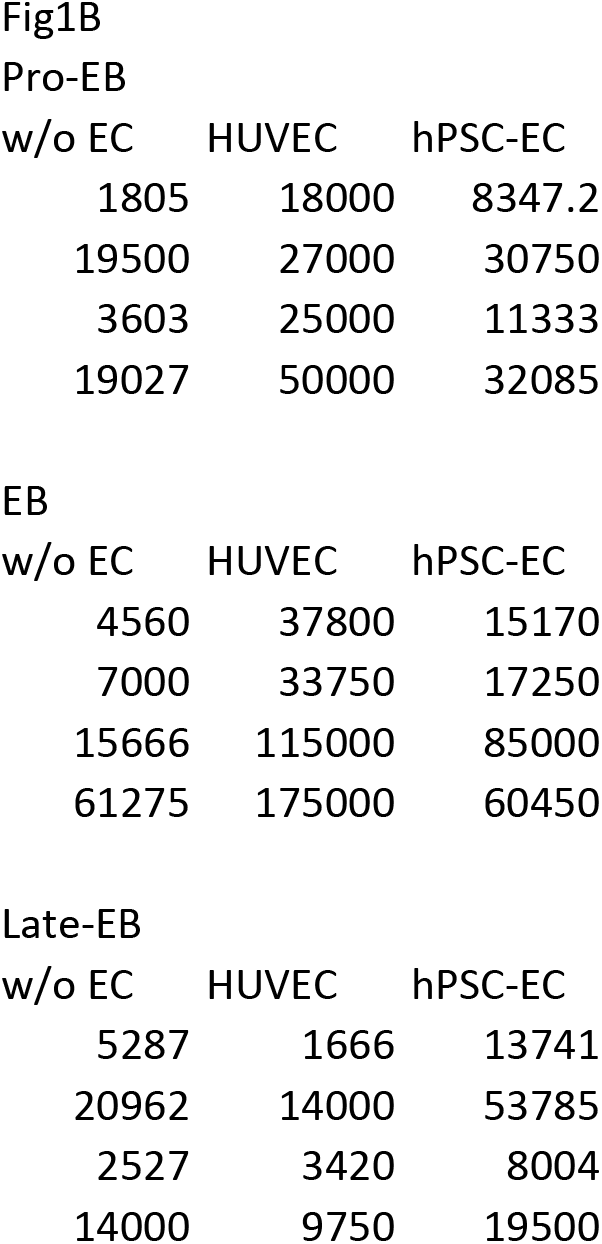
Co-culture with HUVECs amplified erythroid cells from BM-CD34+ cells. (A) Flow cytometry plots of CD71 and GLY -A from BM-CD34+ cells cultured without ECs (left), with HUVECs (middle), and with hPSC-ECs (right). (B) Bar graphs show the number of Pro-EBs (CD71+GLY-A−), EBs (CD71+GLY-A+) and Late-EBs (CD71−GLY-A+). The r ight panel shows a stack of cell numbers for each population. Each dot represents the result from a biologically independent experiment (left three panels). N=4. The data shown as mean ± s.d. * p<0.05 (comparison between w/o EC samples).

**Fig. 2.**
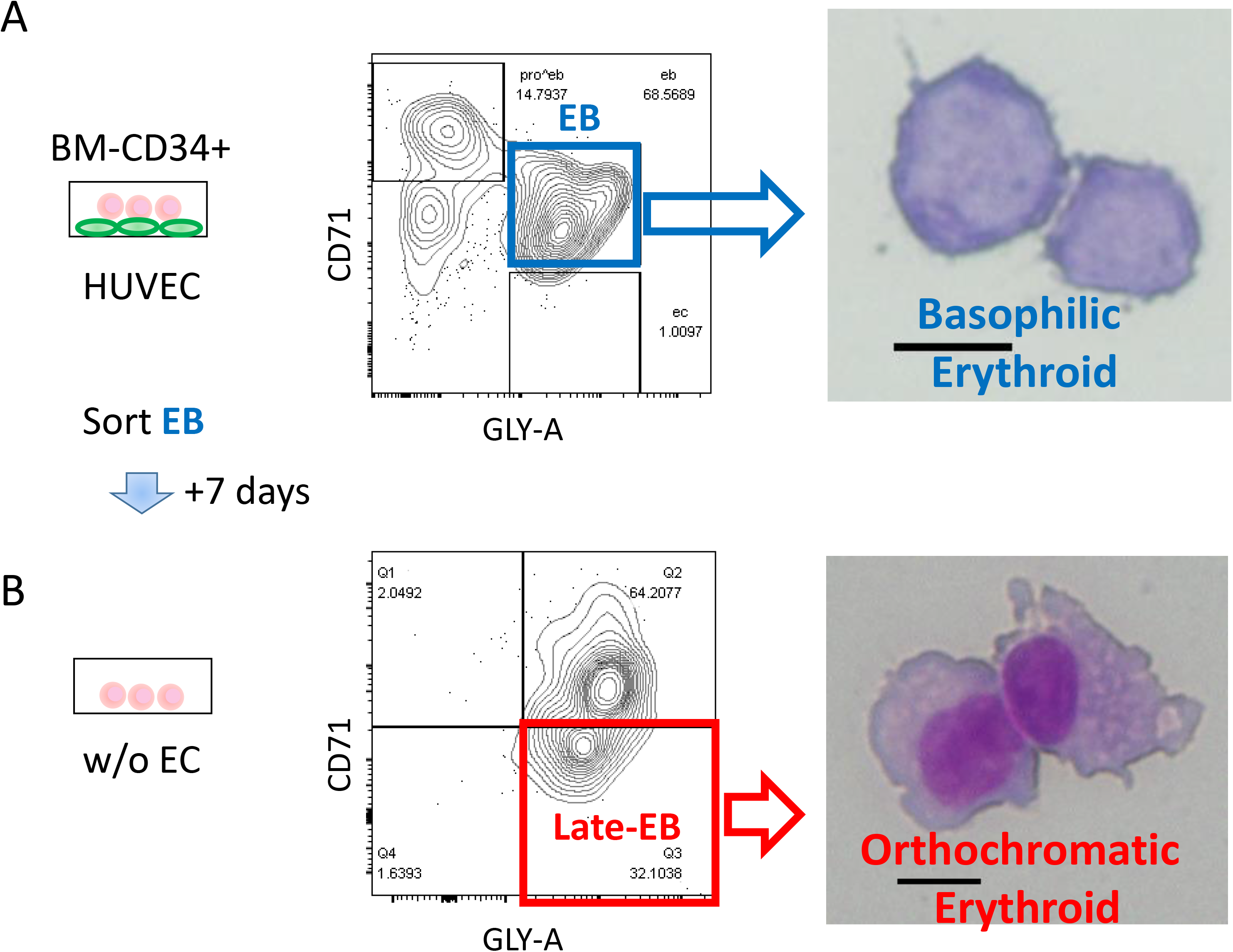
Amplified erythroid cells can differentiate. (A) CD71+ GLY-A+ EBs from BM-CD34+ cells were amplified in the presence of HUVEC s, sorted, and then cultured an additional 7 days. Flow cytometry plots of CD71 and GLY -A from BM-CD34+ cells (middle). Cytospinning shows basophilic erythroids (right). (B) Flow cytometry plots of CD71 and GLY -A from cultured EBs (left). Cytospinning shows orthochromatic erythroids (right). Scale bar = 10 um.

### Angiocrine factors from HUVECs amplify erythroblasts

To assess whether cell-cell interactions are required for the enhanced amplification of erythroid cells, we used a transwell assay, where the media and secreted factors could be exchanged but direct HUVEC contact was prevented. We found an increase in erythroid cells in the transwell setting (fig. 3), suggesting that secreted angiocrine factors are involved in the erythroid amplification. To determine the angiocrine factors produced by HUVECs, we conducted an RNA-seq analysis for the genes that encode secreted factors in HUVECs and hPSC-ECs (fig. 4A). Based on literature search, we classified the secreted factors into three categories (fig. 4B). Erythropoiesis enhancing factors (EEFs) include classical hematopoietic cytokines and morphogens whose roles are validated in erythropoiesis (Paulson, 2011). Recent studies proved that both OP9 (Trakarnsanga, 2018) and macrophages (Lopez-Yrigoyen, 2019) enhance erythropoiesis via secreted factors, which are termed OP9-crines and macrophage-crines respectively in this study. EPO, EGF ligands, and glucocorticoid synthases were included among EEFs, whose expressions were not detected in HUVECs (fig. 4B). Additionally, OP9-crines were not enriched in HUVECs (fig. 4B). On the other hand, NRG1 and IGFBP6, which are included among macrophage-crines, were exclusively expressed in HUVEC s (fig. 4B, table 1). These data demonstrate that HUVECs express and share their profile of angiocrine factors with macrophages.

**Fig. 3.**
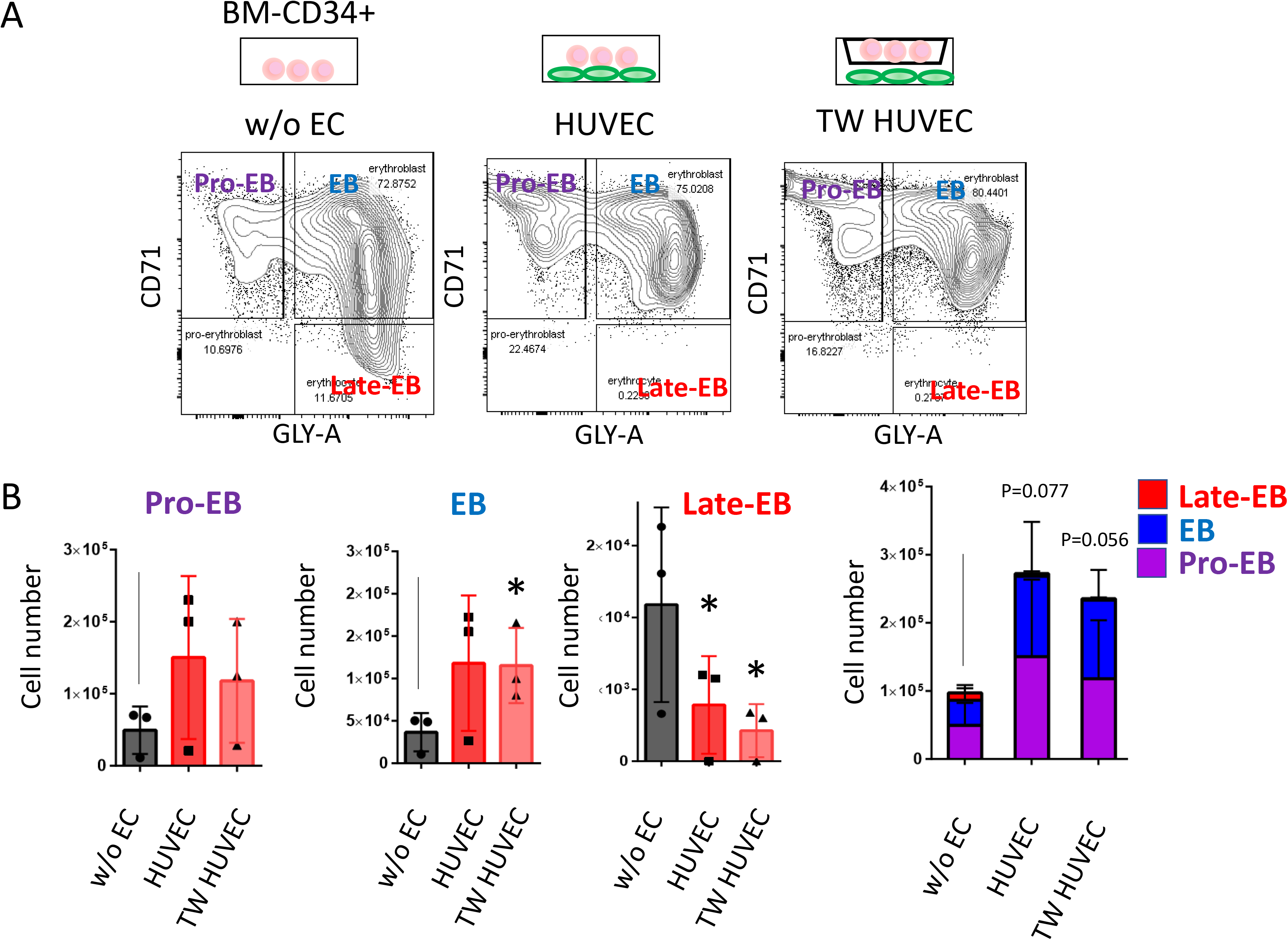
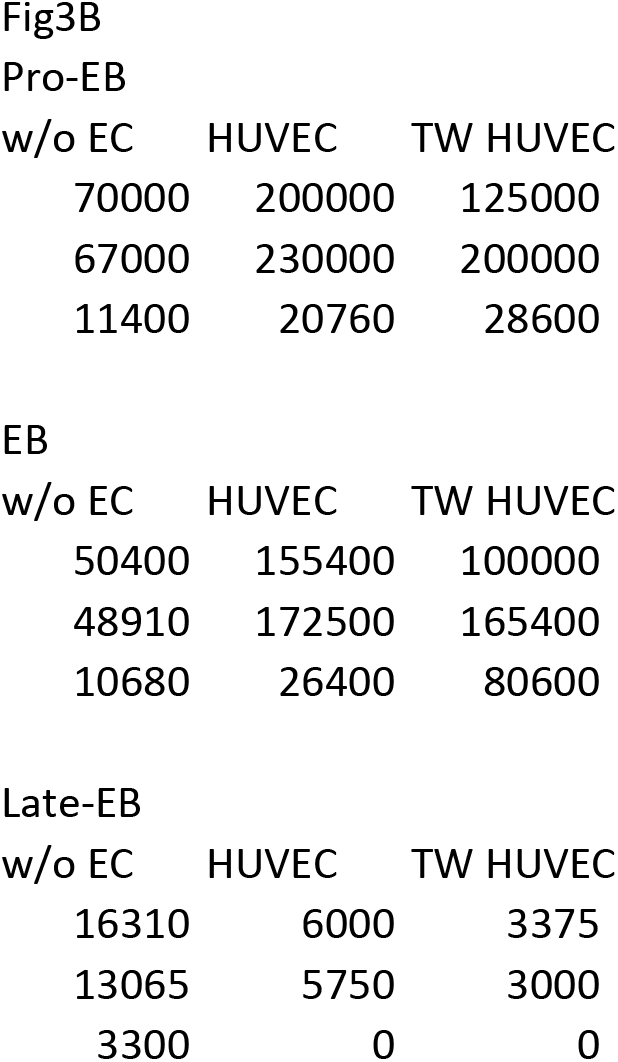
Angiocrine factors from HUVECs amplify erythroid cells from BM-CD34+ cells. (A) Flow cytometry plots of CD71 and GLY -A from BM-CD34+ cells cultured without ECs (left), with HUVECs (middle), and with Transwell (TW)-HUVECs (right). (B) Bar graphs show the number of Pro-EBs (CD71+GLY-A−), EBs (CD71+GLY-A+), and Late-EBs (CD71−GLY-A+). The right panel shows a stack of cell numbers for each population. Each dot represents the result from a biologically independent experiment (left three panels). N=3. The data shown as mean ± s.d. * p<0.05 (comparison between w/o EC samples).

**Fig. 4.**
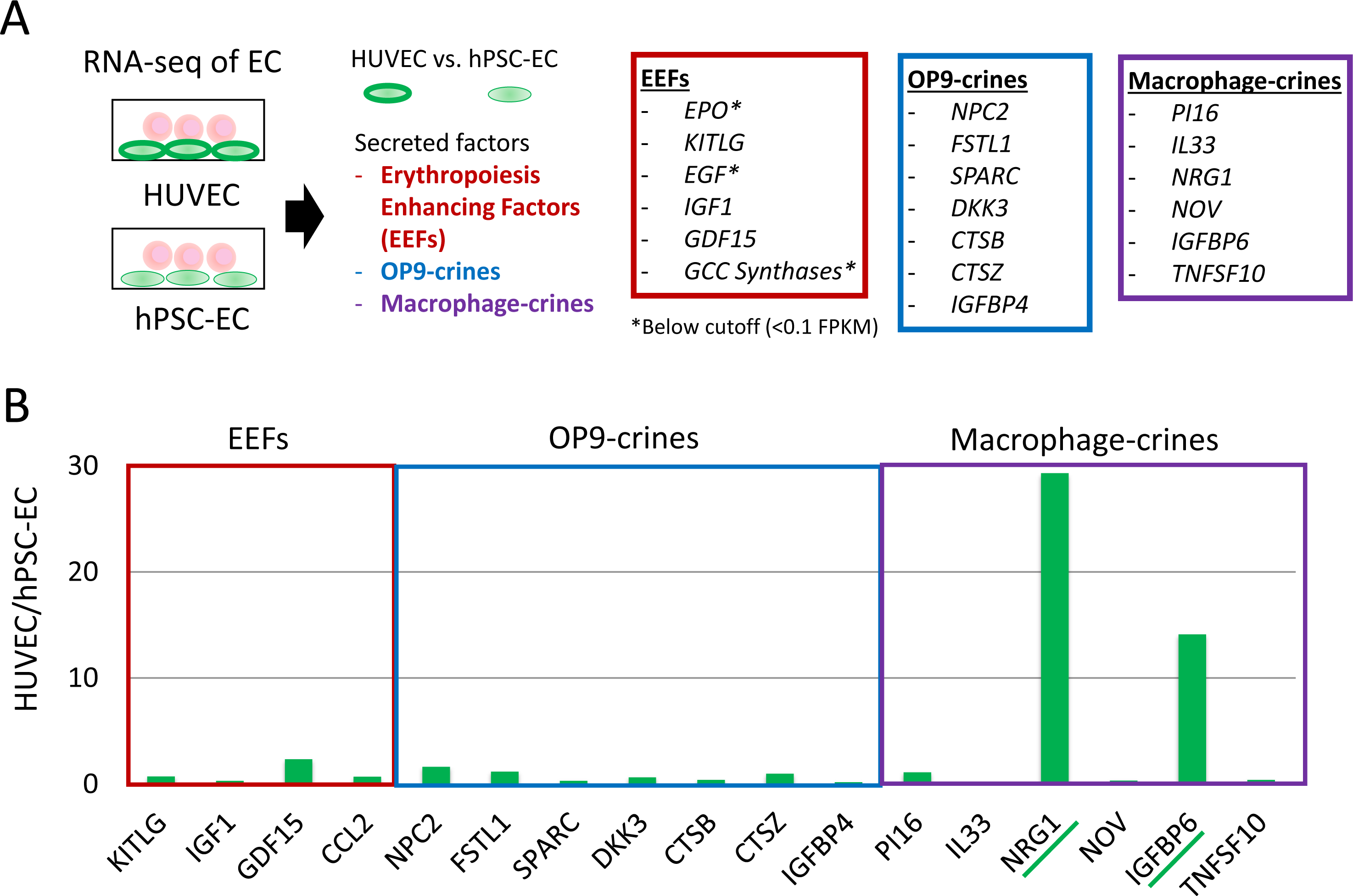
Profiling the angiocrine factors secreted from HUVECs. (A) Bulk RNA-seq analysis of HUVECs and hPSC-ECs was conducted and profiled for erythropoiesis enhancing factors (EEFs) including glucocorticoid (GCC) synthases, OP9-crines and macrophage-crines that are known to promote erythropoiesis (B) Expression of angiocrine factors enriched in HUVECs compared with hPSC-ECs. Refer to Table 1 for the Fragments Per Kilobase of transcript per Million mapped reads (FPKM) of each gene.

**Table.1.**
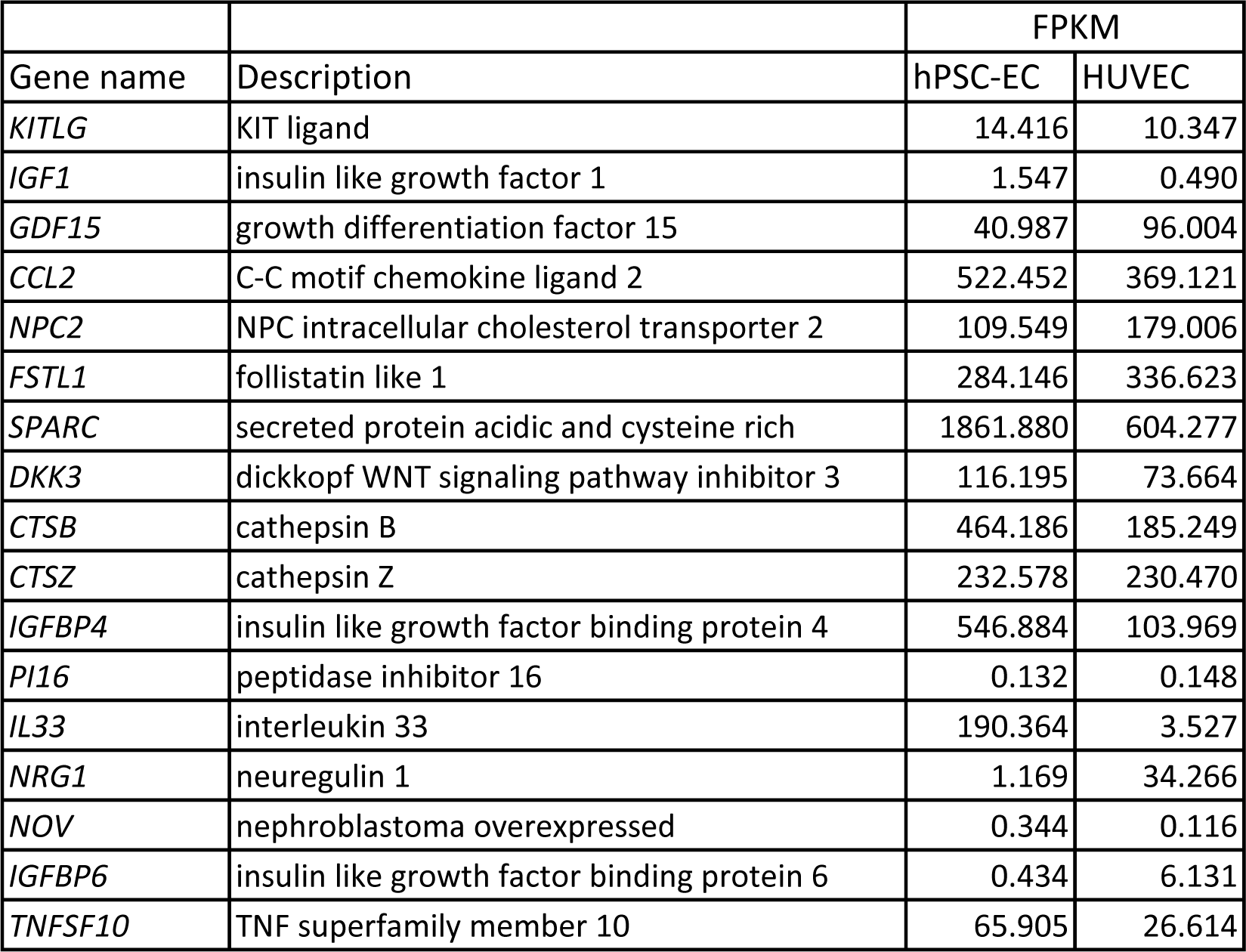
FPKM of erythropoiesis enhancing factors, OP9-crines and macrophage-crines in each EC

### ERK is involved in amplification of erythroid cells

NRG1 and IGFBP6 regulate ERK signaling (fig. 5A; Kataria, 2019; Zinn, 2013). We confirmed that HUVECs activated the ERK signal in erythroid cell line K562 (fig. 5B). Accordingly, we observed that the chemical inhibition of ERK diminished the HUVEC-mediated amplification of erythroid cells (fig. 5C-D). Consistent with this reduced amplification, the inhibition of ERK moved the scatter profile of the GLY-A+ population toward differentiation (fig. 5C). Of note, we made consistent observations in both BM- and cord blood (CB)-CD34+ cells, suggesting the mechanisms of erythroid cell amplification is common between the cell source type: CB from fetus and BM from adult (fig. 5D). These data demonstrate that ERK activation amplifies erythroid cells through angiocrine factors from HUVECs.

**Fig. 5.**
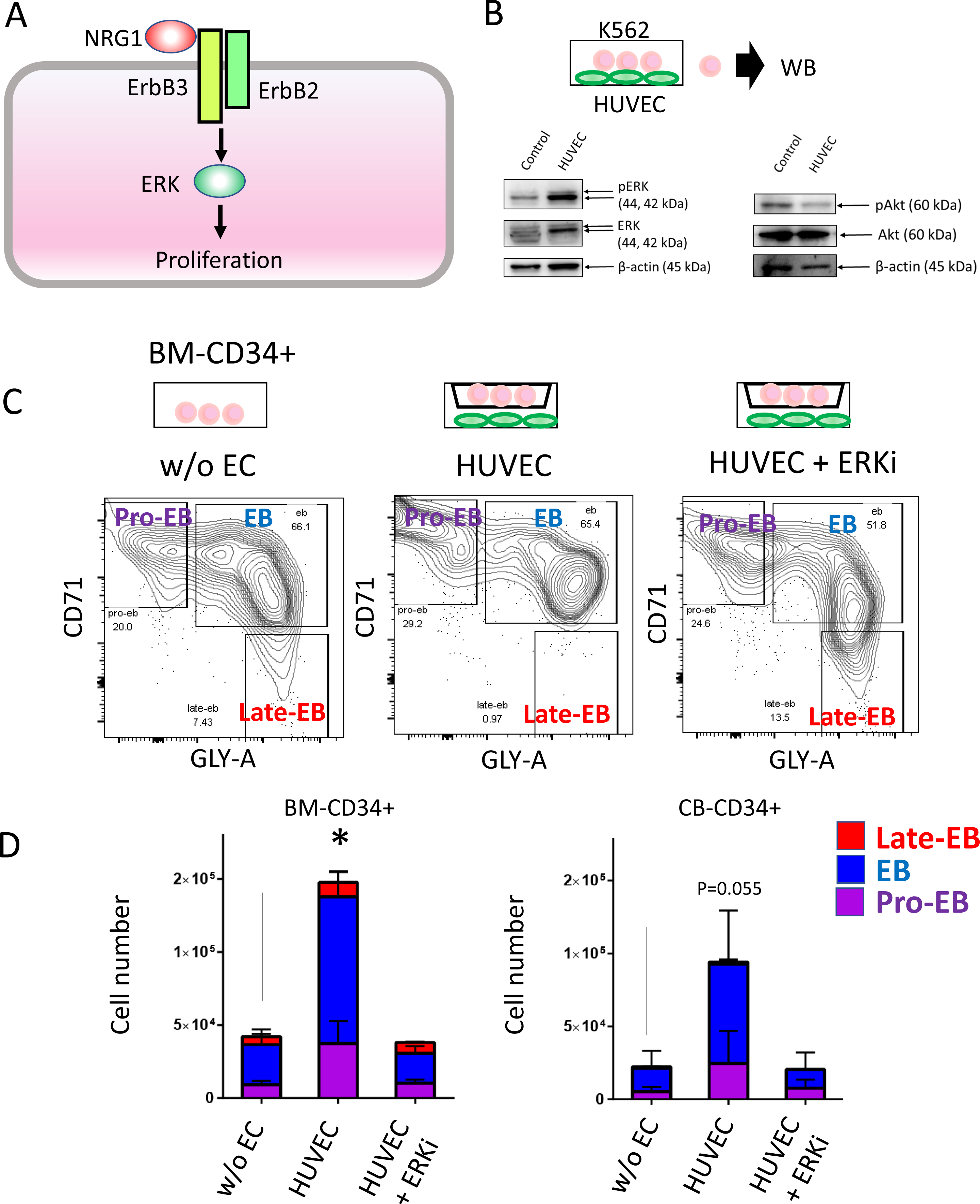
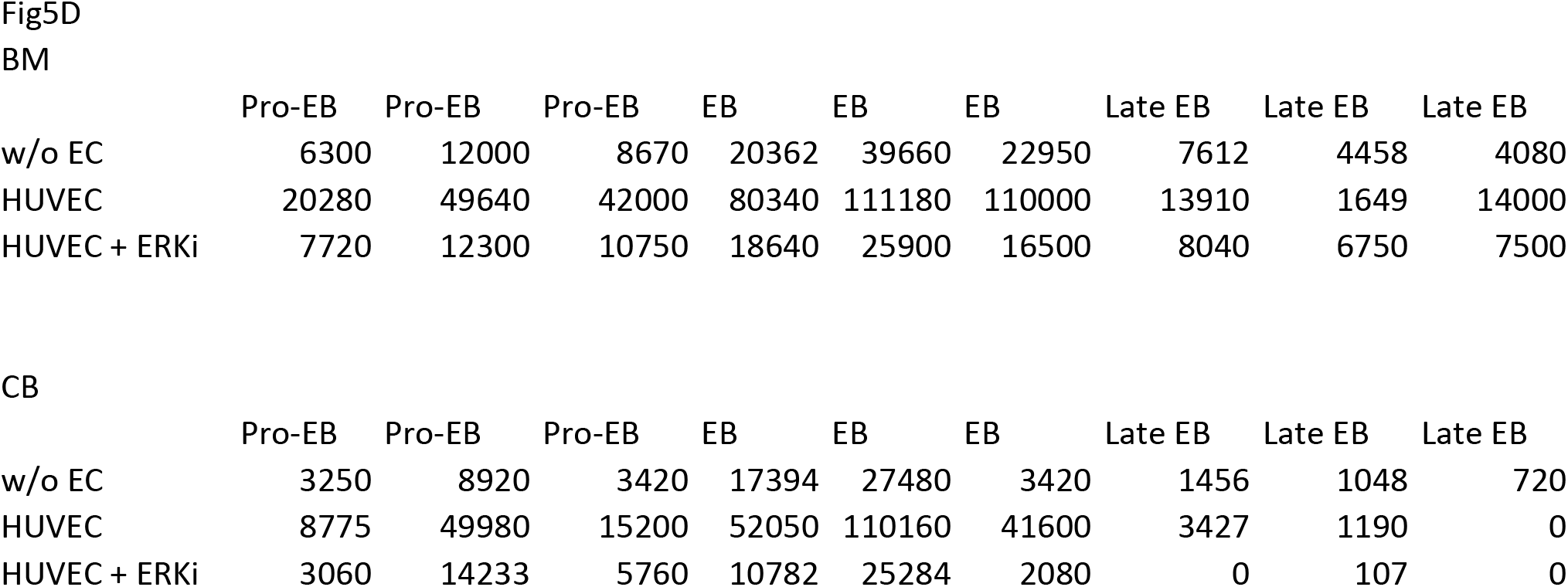
ERK activation amplifies erythroid cells. (A) Scheme of the ERK signal in erythropoiesis. (B) Western blot analysis of the ERK and AKT pathways in K562 cocultured with HUVECs. N=1. (C) Flowcytometry plots of CD71 and GLY -A from BM-CD34+ cells cultured without ECs (left), with HUVECs (middle), and HUVECs + ERK inhibitor (right). (D) A stack of cell numbers for Pro-EBs, EBs, and Late-EBs from the experiments from BM-CD34+ and CB-CD34+ cells. N=3. The data shown as mean ± s.d. * p<0.05 (comparison between w/o EC samples).

## Discussion

Elucidating a manner to amplify erythroid cells is expected to achieve a clinically relevant number of red blood cells (Koury, 2016). Several studies have shown that HUVECs proliferate hematopoietic stem and progenitor cells (Yildirim, 2005), but the understanding of their role in erythropoiesis remains incomplete. Our work demonstrated that angiocrine factors secreted by HUVECs amplify erythroid cells, suggesting a possible mechanism to produce red blood cells *in vitro*.

ERK signaling mediates erythroid commitment of hematopoietic progenitor cells at the initial phase of erythropoiesis, but its role in the subsequent phases erythropoiesis is not clear (Guihard, 2010). We demonstrated that ERK signaling amplifies erythroid cells through angiocrine factors from HUVECs. Instead of other erythropoiesis factors, such as EPO (Kuhrt, 2015), SCF (Comazzetto, 2019) or glucocorticoids (Narla, 2011), HUVECs produce angiocrine factors in common with those from macrophages to enhance erythropoiesis (Lopez-Yrigoyen, 2019; Seu, 2017). One study showed the contribution of cell-cell contacts between murine hemangioma-derived ECs and erythropoiesis. However, we did not find these contacts to be a factor with HUVECs. (Ohneda, 1997). To conclude, our findings suggest that angiocrine factors could enhance erythropoiesis from either donor-derived or hPSC-derived hematopoietic progenitor cells.

## ACKNOWLEDGEMENTS

We are grateful to Alina Li and Takafumi Mano for their technical assistance and Dr. Daisuke Okuzaki for the RNA-seq analysis. We would also like to thank Ms. Harumi Watanabe for providing administrative assistance and Dr. Peter Karagiannis for reading and editing the paper.

This work was supported by the Core Center for iPS Cell Research of Research Center Network for Realization of Regenerative Medicine from the Japan Agency for Medical Research and Development (AMED) [M.K.S.], the Program for Intractable Diseases Research utilizing Disease-specific iPS cells of AMED (17935423) [M.K.S.], AMED under Grant No. JP18gm5810008 [Y.-s.T.], JSPS KAKENHI Grant No. JP17H02082 [Y.-s.T.], the Kyoto University Hakubi Project, and the Center for Innovation program of Japan Science and Technology Agency (JST) [R.O. and M.K.S.]. R.S. is a recipient of Early Career KAKENHI, iPS Academia Japan and Sen-shin Medical Research Foundation (SMRF) fellowships. The authors declare no conflicts of interest.

## AUTHOR CONTRIBUTIONS

R.S. and R.O. designed the study, conducted the experiments, interpreted the data and wrote the manuscript. C.M., E.S., T.S. T.N., and Y-s..T. conducted the experiments. M.K.S. supervised the study. R.S., R.O., Y-s.T., A.N., and M.K.S. commented on and wrote the paper.

## Disclosures

The authors declare they have no competing financial interests.

## Methods

### Contact for Reagent and Resource Sharing

Further information and requests for resources and reagents should be directed to and will be fulfilled by the Lead Contact, Ryohichi Sugimura.

### Experimental Model and Subject Details

#### Cell lines

All the experiments of this study were performed with 409B2 iPSC and CBA11 iPSC lines (Ohta, 2019). K562 was obtained through RIKEN Bioresource Center. Human BM-CD34+ cells were purchased from Lonza (Tissue Acquisition Number: 35843, 35845, 32423, 34781, 30968). Human CB-CD34+ cells were purchased from Stemexpress (mixed donor, CB34P3401C). HUVECs were either purchased from Angio-Proteomie (GFP-HUVEC, cAP-01001GFP) or Lonza (issue Acquisition Number: 29000).

### Accession number of RNA-sequencing

GSE138104 https://www.ncbi.nlm.nih.gov/geo/query/acc.cgi?acc=GSE138104

## Method details

### hPSC culture

The maintenance of hPSCs was done using iMatrix-511 (Matrixome) in mTeSR1 media (STEMCELL Technologies). Media were changed every other day, and the cells were passaged as single cells every 7 days using TrypLE Express (Life technologies).

### Endothelial differentiation

hPSC spheroids were formed as described previously (Ohta, 2019). hPSC spheroids were suspended in mTeSR1 (StemCell Technologies) containing 1.25 μg/mL iMatrix-511 (Matrixome) and subsequently plated onto non-coated culture plates. After three days, the medium was replaced with Essential 8 (Life technologies) containing 4 μM CHIR99021 (WAKO), 80 ng/mL BMP4 (R&D systems), and 80 ng/mL VEGF (R&D systems). After two more days, the cells were dissociated to the single-cell level with TrypLE Express (Life technologies) for 20 minutes at 37° and subsequently plated onto an iMatrix-411 (Matrixome)-coated plate in Stempro34-SFM (Life technologies) containing 80 ng/mL VEGF. After four more days of culture on iMatrix-411, the cells were passaged onto a Type I collagen-coated plate at a density of 10,000/cm^2^ in EGM-2 containing 25 ng/mL VEGF and cultured for 9 days with passage every four days.

### Co-culture

20,000 BM or CB CD34+ cells were cultured with 60,000 HUVECs or hPSC-derived ECs seeded on 50 μg/mL of fibronectin-coated plates in alpha-MEM medium supplemented with 15% FBS, 1% 5 insulin-transferrin, ng/mL IL-7, 10 ng/mL FLT-3L, 50 μg/mL L-ascorbic acid, and 1% L-Glut/Pen/Strep. The cells were co-cultured for 7 days. For the transwell analysis, we used a 0.4 μM pore transwell (MCHT12H48 Millipore) and seeded CD34+ cells on the transwell apparatus. ERK inhibitor FR180204 (10 μM) was added to culture where described in the text.

### Cytospin

5,000 FACS-sorted CD71+GLY-A+ or CD71−GLY-A+ cells were cytospun onto slides (500 r.p.m. for 5 min), air dried, and stained with MayGrunwald and Giemsa stains, washed with water, air dried, and mounted, followed by examination by light microscopy.

### Erythroid differentiation of K562

K562 cells were maintained in Ham F12 with 10% FBS. For erythroid differentiation, sodium butyrate was added to the medium (1 mM) and cultured for 7 days.

### Antibody

Anti-pERK (#4370), anti-ERK (#4695) and anti-beta-actin (#4970) were purchased from Cell Signaling Technology.

### Western blot

Equal amounts of protein extracted from K562 cells were subjected to SDS-PAGE in Tris-Glycine buffer and transferred to PVDF membranes. The membrane was blocked with 5% Skim milk in Tris-buffered saline containing 0.1% Tween-20 (TBS-T) for 1 hour at room temperature and probed with the appropriate primary antibody (1:1,000, anti-pERK, ERK or beta-actin antibody) overnight at room temperature. After washing with TBS-T, the membrane was incubated with the appropriate secondary antibody (anti-rabbit IgG HRP-linked antibody (1:1,000, cell signaling)) for 1 hour at room temperature. After washing with TBS-T, the membrane was incubated with West Femto super signal reagent (Thermo scientific), and the specific proteins were visualized with LAS-4000 (GE healthcare).

### Flow cytometry

Cells grown in culture or harvested from animal tissues were stained with 4:200-1:200 dilution of each antibody for at least 30 min on ice in the dark with the following antibodies; CD71-PE and GLY-A-PECY7. Unless specifically indicated, all the antibodies used are against human cells. Acquisitions were done on a BD FACSAria II cell sorter or BD LSRFortessa cytometer. Sorting was performed on a BDFACS Aria II cell sorter. Flow cytometry data were analyzed using FlowJo V.10.

### Bulk RNA-seq

Sequencing was performed on an Illumina HiSeq 2500 platform in a 75 -base single-end mode. Illumina Casava1.8.2 software was used for basecalling. Sequenced reads were mapped to the human reference genome sequences (hg19) using TopHat v2.0.13 in combination with Bowtie2 ver. 2.2.3 and SAMtools ver. 0.1.19. The fragments per kilobase of exon per million mapped fragments (FPKMs) was calculated using Cufflinks version 2.2.1.

### Statistics and source data

Statistical analyses were done with t-test. We used Microsoft Excel for calculations and expressed the results as the means ± s.d. The source data for each graph is available in the supplementary tables.

## Notes

https://www.ncbi.nlm.nih.gov/geo/query/acc.cgi?acc=GSE138104

## REFERENCES

An X, Schulz VP, Mohandas N, Gallagher PG. Curr Opin Hematol. 2015, 22, 3.

Baron MH, Isern J, Fraser ST. Blood, 2012, 119, 21.

Batta K, Menegatti S, Garcia-Alegria E, Florkowska M, Lacaud G, Kouskoff V. Stem Cells Transl Med, 2016, 5, 10.

Butler JM, Nolan DJ, Vertes EL, Varnum-Finney B, Kobayashi H, Hooper AT, Seandel M, Shido K, White IA, Kobayashi M, Witte L, May C, Shawber C, Kimura Y, Kitajewski J, Rosenwaks Z, Bernstein ID, Rafii S. Cell Stem Cell, 2010, 6, 3.

Chung SS, Park CY. Blood Adv, 2017, 1, 26.

Doulatov S, Daley GQ. Science, 2013, 342, 6159.

Eblen ST, Adv Cancer Res. 2018, 138.

Fang S, Nurmi H, Heinolainen K, Chen S, Salminen E, Saharinen P, Mikkola HK, Alitalo K. Blood, 2016, 128, 5.

Grasman JM, Kaplan DL. Sci Rep. 2017, 7, 1.

Kataria H, Alizadeh A, Karimi-Abdolrezaee S. Prog Neurobiol. 2019, 180.

Kinney MA, Vo LT, Frame JM, Barragan J, Conway AJ, Li S, Wong KK, Collins JJ, Cahan P, North TE, Lauffenburger DA, Daley GQ. Nat Biotechnol, 2019, 37, 7.

Comazzetto S, Murphy MM, Berto S, Jeffery E, Zhao Z, Morrison SJ, Cell Stem Cell, 2019, 24, 3.

Guihard S, Clay D, Cocault L, Saulnier N, Opolon P, Souyri M, Pagès G, Pouysségur J, Porteu F, Gaudry M, Blood, 2010, 115, 18.

Koury MJ. Exp Hematol. 2016, 4.

Kuhrt D, Wojchowski DM. Blood, 2015, 125, 23.

Li H, Natarajan A, Ezike J, Barrasa MI, Le Y, Feder ZA, Yang H, Ma C, Markoulaki S, Lodish HF. Dev Cell, 2019, 49, 1.

Lopez-Yrigoyen M, Yang CT, Fidanza A, Cassetta L, Taylor AH, McCahill A, Sellink E, von Lindern M, van den Akker E, Mountford JC, Pollard JW, Forrester LM. Nat Commun. 2019, 10, 1.

Morrison SJ, Scadden DT. Nature, 2014, 505, 7483.

Nandakumar SK, Ulirsch JC, Sankaran VG. Br J Haematol. 2016, 173, 2.

Narla A, Dutt S, McAuley JR, Al-Shahrour F, Hurst S, McConkey M, Neuberg D, Ebert BL. Blood, 2011, 118, 8.

Ohneda O, Bautch VL, Br J Haematol. 1997, 98, 4.

Ohta R, Sugimura R, Niwa A, Saito MK. J Vis Exp. 2019, 148.

Orkin SH, Zon LI. Cell, 2008, 132, 4.

Paulson RF, Shi L, Wu DC. Curr Opin Hematol. 2011, 18, 3.

Pimkin M, Kossenkov AV, Mishra T, Morrissey CS, Wu W, Keller CA, Blobel GA, Lee D, Beer MA, Hardison RC, Weiss MJ. Genome Res. 2014, 24, 12.

Pinho S, Frenette PS. Nat Rev Mol Cell Biol. 2019, 20, 5.

Rezaei M, Martins Cavaco AC, Seebach J, Niland S, Zimmermann J, Hanschmann EM, Hallmann R, Schillers H, Eble JA. J Immunol. 2019, 202, 5.

Seu KG, Papoin J, Fessler R, Hom J, Huang G, Mohandas N, Blanc L, Kalfa TA. Front Immunol. 2017, 8, 1140.

Smalley I, Smalley KSM. Cancer Discov. 2018, 8, 2.

Trakarnsanga K, Wilson MC, Heesom KJ, Andrienko TN, Srisawat C, Frayne J. Sci Rep. 2018, 8, 1.

Van Handel B, Prashad SL, Hassanzadeh-Kiabi N, Huang A, Magnusson M, Atanassova B, Chen A, Hamalainen EI, Mikkola HK. Blood, 2010, 116, 17.

Wei Q, Boulais PE, Zhang D, Pinho S, Tanaka M, Frenette PS. Blood, 2019, 133, 11.

Yildirim S, Boehmler AM, Kanz L, Möhle R. Bone Marrow Transplant, 2005, 36, 1.

Zinn RL, Gardner EE, Marchionni L, Murphy SC, Dobromilskaya I, Hann CL, Rudin CM. Mol Cancer Ther. 2013, 12, 6.

Ziyad S, Riordan JD, Cavanaugh AM, Su T, Hernandez GE, Hilfenhaus G, Morselli M, Huynh K, Wang K, Chen JN, Dupuy AJ, Iruela -Arispe ML. Cell Rep. 2018, 22, 5.

